# MicroRNA Regulatory Targets Related to VO_2_peak Decline in Older Adult Participants of the Study of Muscle, Mobility and Aging (SOMMA)

**DOI:** 10.64898/2026.01.19.700188

**Authors:** Genesio M. Karere, Fang-Chi Hsu, Russell T. Hepple, Paul M. Coen, Steve R. Cummings, Anne B. Newman, Nancy W. Glynn, Lauren M. Sparks, Nancy E. Lane, Albert G. Hayward, Jianzhao Xu, Nathan Wagner, Ge Li, Jeanne Chan, Laura A. Cox, Stephen B. Kritchevsky

## Abstract

Skeletal muscle aging (sarcopenia) is associated with reduced peak oxygen consumption (VO peak) during exercise, a key determinant of physical function and overall health. However, the molecular mechanisms linking muscle aging to low VO peak remain poorly understood. We aimed to identify miRNA signatures and miRNA–gene regulatory networks associated with VO peak in older adults. Using small RNA and mRNA sequencing, we analyzed skeletal muscle from 72 SOMMA participants (70–79 years old) with low or high VO peak (n = 18/group) and from 36 participants spanning the full VO peak spectrum. Differential expression was assessed using LIMMA, with pathway and network analyses performed using Ingenuity Pathway Analysis (IPA) and Weighted Gene Co-expression Network Analysis (WGCNA). We detected 1,408 miRNAs and 16,210 genes; among these, 14 miRNAs and 2,018 genes were differentially expressed (FDR < 0.05). The 14 miRNAs regulated 142 genes, and expression of 10 miRNAs inversely correlated with 50 genes enriched in mitochondrial, sirtuin-1, and nitric oxide signaling pathways. Regression analyses identified 21 miRNAs and 1,744 genes significantly correlated with VO peak after adjusting for age and sex. WGCNA revealed 10 co-expression modules associated with VO peak, with the cyan module showing the strongest correlation and enrichment for nitric oxide signaling genes. These findings highlight novel miRNA-mediated molecular pathways potentially contributing to low VO peak and skeletal muscle aging in older adults. Future studies will further investigate these miRNA–gene interactions to uncover therapeutic targets for preserving muscle function with age.

## INTRODUCTION

Skeletal muscle aging is characterized by progressive loss of mass and functional decline (sarcopenia), a major contributor to impaired mobility in older adults (1,2). In the United States, sarcopenia affects an estimated 36% of individuals with a mean age of 70 years (3), with prevalence of 4.6% in men and 7.9% in women (4). Sarcopenia is closely associated with reduced cardiorespiratory fitness (CRF), most commonly assessed by peak oxygen consumption (VO peak) during cardiopulmonary exercise testing (5). VO peak correlates with several clinically relevant outcomes, including 400-m walk time (6), gait speed (7), mitochondrial energetics (8), life-space mobility (5,9), fatigability (10–12), and leg power (6,7), making it a robust indicator of overall physical fitness. However, substantial interindividual variation in VO peak exists among older adults, and the molecular mechanisms underlying this variability remain poorly understood. Defining these mechanisms is essential for identifying skeletal muscle pathways that contribute to VO peak differences and for developing targeted interventions to improve fitness in aging populations.

We posit that skeletal muscle microRNAs (miRNAs) and miRNA gene targets correlate with and are enriched in pathways related to VO_2_peak variation in older adults. miRNAs are small non-coding RNAs that post-transcriptionally regulate gene expression, altering cellular protein abundance and processes including cell proliferation, differentiation, apoptosis, and metabolic homeostasis at cellular and *in-vivo* levels (13–16). The functional role of miRNAs in influencing cellular protein abundance is relevant to the pathogenesis of sarcopenia, as is caused primarily by dysregulation of protein homeostasis in muscle cells due to imbalance of protein synthesis and degradation. Previous studies have demonstrated that the dysregulation of miRNA expression with age contributes to muscle sarcopenia by regulating the PI3-AKT/FOXO pathway that is implicated in ubiquitin-proteosome system, myogenesis, cell apoptosis and mitochondria dysfunction (17). Other studies have shown that miRNAs and mRNAs (genes) are responsive to exercise interventions (18–22) and regulate skeletal muscle development (23–25). However, the mechanisms in which skeletal muscle miRNA-mediate VO_2_peak variations are not fully understood. To our knowledge, no prior studies have leveraged multiomic approaches to understand the skeletal muscle miRNA-mediated molecular mechanisms related to VO_2_peak variation in older adults. However, cumulative evidence demonstrates the role of miRNAs in skeletal muscle aging. A study in a mouse model identified 57 skeletal muscle miRNAs that were differentially expressed between young (n=24 mice, 12 months old) and old (n=24 mice, 24 months old) using TaqMan miRNA array (26). Using next generation sequencing, another study identified skeletal muscle expressed miRNAs that differed between young and old mice (age was not specified in this study) (27). In rhesus monkeys, the expression of 35 miRNAs differed between young (6 years old, n=4) and old (26.8 years old, n=4) (28). In human studies, a comparison of skeletal muscle miRNA expression profiles between young (31 years, n=19) and old (73 years, n=17) using miRNA array identified 18 miRNAs that differ between groups including miR-181a/b (13). These studies in animal models and in humans have demonstrated a potential role for miRNAs in skeletal muscle aging. Other studies interrogated the effects of combined aging and exercise on miRNA expression in skeletal muscles (13, 29). Together, these studies confirm the regulatory role of miRNAs in skeletal muscle aging. However, limitations of these studies include the small sample sizes analyzed and not directly addressing VO_2_peak variation in older adults. In addition, the employment of targeted approaches may have hindered the discovery of novel findings. In contrast, we used an unbiased, untargeted RNA sequencing approaches (including small RNAseq) combined with a multiomic analytical approach to identify all differentially expressed miRNAs and miRNA target genes that may be responsible for VO_2_peak variation in older adults and in a larger sample size.

The goal of our study was to identify skeletal muscle miRNA signatures, miRNA regulatory targets, and dysregulated pathways that contribute to low VO_2_peak in older adults. To accomplish this goal, we leveraged baseline skeletal muscle samples from a cohort of the participants of the Study of Muscle, Mobility and Aging (SOMMA) (30). Recruitment and clinical testing of the participants of SOMMA were conducted at the University of Pittsburgh (Pittsburgh, PA) and Wake Forest University School of Medicine (Winston-Salem, NC), with overall objective of understanding the impact of aging skeletal muscle on mobility. Inclusion criteria included body mass index (BMI) of 18–40 kg/m^2^, ≥70 years old at enrollment, and were willing to complete magnetic resonance imaging and a muscle tissue biopsy. Participants were excluded if unable to walk one quarter of a mile or climb a flight of stairs or having an active malignancy or had an advanced chronic disease such as heart failure, renal failure on dialysis, Parkinson’s disease, or dementia (30). We reveal a novel signature of skeletal muscle miRNAs that correlates with VO_2_peak variation, as well as a miRNA-gene molecular network and enriched pathways related to VO_2_peak. Our findings highlight critical insights into the molecular changes that drive low VO_2_peak and potential therapeutic targets for intervention to slow skeletal muscle aging to reduce immobility and mortality in older individuals.

## METHODS

### Study Design

The study design included a discovery cohort and a validation cohort. The discovery cohort was comprised of SOMMA participants selected for extreme values of VO_2_peak to capture the greatest extent of genetic variation in SOMMA (N=18 with very low VO_2_peak (12.8±1.1 mL/kg/min), and 18 with very high VO_2_peak (33.4±4.0 mL/kg/min)). The validation cohort included the 36 randomly selected SOMMA participants spanning the entire spectrum of VO_2_peak measures from low to high VO_2_peak (12.7 to 30.6 mL/kg/min) to identify miRNAs that relate to VO_2_peak.

### VO_2_peak and Skeletal Muscle Sample Collections

VO_2_peak was assessed during cardiopulmonary exercise testing as previously described (31). Briefly, VO_2_peak was quantified as the highest volume of oxygen consumption averaged over 30-second intervals with progression of exercise intensity in a treadmill, calculated using BreezeSuite metabolic cart software (MGC Diagnostics, St. Paul, MN) and adjusted for participant weight. Skeletal muscle biopsies were collected and processed from the middle region of the musculus vastus lateralis under local anesthesia using a Bergstrom canula with suction as described previously (8, 32).

### Small RNA Sequencing

We identified miRNAs expressed in skeletal muscles by performing small RNAseq and sequence analyses on RNA isolated from flash frozen muscle biopsy samples as described in our previous studies (33, 34). Briefly, we used 50ng of total RNA to generate cDNA libraries using NextFlex Small RNASeq Kit v4 (Revvity) and SciClon Workstation Robotics (Revvity). cDNA quality was assessed using TapeStation (Agilent). We performed small RNAseq using Illumina NovaSeq 6000 instrument and reagents. We used mirDeep2 pipeline (35) and Partek Flow (Illumina) to analyze fastq-formatted sequence reads with a Phred quality score of 30 or greater, an inferred base call accuracy of 99.9%, to identify miRNAs and assess their expression levels (read counts). miRNA read counts were normalized holistically using reads per million mapped to a miRNA.

In addition, data on skeletal muscle-expressed mRNAs (genes) were obtained from the SOMMA Coordinating Center. Quality control and normalization of the data were performed as described previously (32). Importantly, expressed miRNAs and genes were identified from the same skeletal muscle samples.

### MiRNA Target Identification, MiRNA-gene Networks, and Pathway Analysis

We performed these steps as previously described (33). Briefly, lists of differentially expressed miRNAs and genes were imported to Ingenuity Pathway Analysis (IPA), and we used the miRNA Target Filter tool in IPA to integrate the miRNA and gene expression datasets. Putative miRNA target genes were identified using TargetScan prediction algorithm in IPA as well as TarBase and miRecords databases, and results were filtered to only include experimentally validated interactions between miRNAs and mRNA targets. We then analyzed miRNA target genes using Core Analysis tool in IPA to identify significant pathways enriched with target genes.

### Weighted Gene Co-expression Analysis (WGCNA)

We used WGCNA in R package to analyze target gene datasets and to identify gene modules and genes in modules that correlate with VO_2_peak. WGCNA is a systems biology method used to describe the correlation patterns among genes, enabling the identification of modules, clusters of highly co-expressed genes, that may reflect shared biological functions or regulatory mechanisms. Co-expression networks were constructed using the ‘blockwiseModules’ function. Genes in the same module are significantly interconnected and indicate a common function class. Correlation between a module and VO_2_peak was visualized using the ‘labeledHeatmap’ of the WGCNA package. Genes in the modules that exhibit the most significant correlation with VO_2_peak were imported to IPA for core analyses to identify significant enriched pathways. Modules were considered significant if the correlation coefficient (r) was ≥ 0.30 and False Discovery Rate (FDR) adjusted p-value < 0.05, indicating a moderate-to-strong and statistically significant relationship with VO peak.

### Data Analysis

We performed differential miRNA expression (e.g., comparing low vs. high VO_2_peak) and regression analyses (e.g., modeling VO_2_peak) using linear models for microarray data (LIMMA) (36), a linear model-based R/Bioconductor analysis software with empirical Bayes variance shrinkage to improve variance estimation. The covariates included age and sex where applicable. Normalized miRNA sequence read counts were log_2_ transformed. Additionally, the penalized regression analysis using least absolute shrinkage and selection operator (LASSO) (37), a machine learning approach implemented in the R *glmnet* package, was used to identify a subset of miRNAs most predictive of VO_2_peak. Model tuning and penalty parameter selection were guided by cross-validation, ensuring that the model’s predictive performance was optimized while minimizing overfitting. In all analyses, the results were considered significant if the FDR adjusted p-value < 0.05.

## RESULTS

### Study Participants

The overall characteristics of the study participants as well as specific characteristics related to individuals with low or high VO_2_peak and randomly selected samples are shown in Table 1.

**Table 1:**
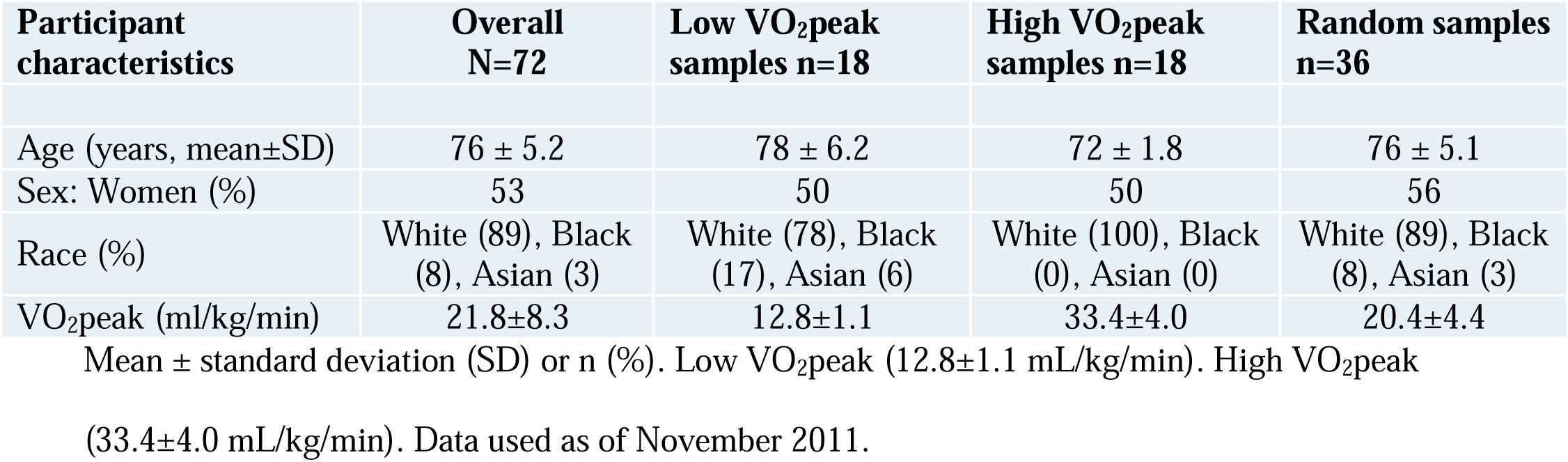
Characteristics of the participants of the SOMMA cohort analyzed in our study.

| Participant characteristics | Overall N=72 | Low VO<sub>2</sub>peak samples n=18 | High VO<sub>2</sub>peak samples n=18 | Random samples n=36 |
| --- | --- | --- | --- | --- |
| Age (years, mean±SD) | 76 ± 5.2 | 78 ± 6.2 | 72 ± 1.8 | 76 ± 5.1 |
| Sex: Women (%) | 53 | 50 | 50 | 56 |
| Race (%) | White (89), Black (8), Asian (3) | White (78), Black (17), Asian (6) | White (100), Black (0), Asian (0) | White (89), Black (8), Asian (3) |
| VO<sub>2</sub>peak (ml/kg/min) | 21.8±8.3 | 12.8±1.1 | 33.4±4.0 | 20.4±4.4 |

Mean ± standard deviation (SD) or n (%). Low VO<sub>2</sub>peak (12.8±1.1 mL/kg/min). High VO<sub>2</sub>peak
(33.4±4.0 mL/kg/min). Data used as of November 2011.

### MiRNAs and Genes Differentially Expressed Between Low and High VO_2_peak Individuals

A total of 1,408 miRNAs and 16,210 genes expressed in skeletal muscle samples, of which 68 miRNAs and 4,748 genes differed between individuals with low and high VO_2_peak based on nominal p-values (**Tables S1** and **S2**). After p-value adjustment, 14 miRNAs and 2,018 genes remained significant (**Table 2** and **Table S2**).

**Table 2:** Differentially expressed skeletal muscle miRNAs between low and high VO_2_peak participants.

| miRNA ID | Log Fold-change | p-value | FDR<0.05 |
| --- | --- | --- | --- |
| hsa-miR-515-5p | 1.6 | 0.00002 | 0.00607 |
| hsa-miR-520g-3p | 1.4 | 0.00002 | 0.00607 |
| hsa-miR-519c-3p | 1.4 | 0.00005 | 0.01122 |
| hsa-miR-517a-3p | 1.2 | 0.00012 | 0.01987 |
| hsa-miR-503-5p | -1.4 | 0.00014 | 0.01987 |
| hsa-miR-518b | 1.5 | 0.00022 | 0.02599 |
| hsa-miR-517b-3p | 1.2 | 0.00043 | 0.04135 |
| hsa-miR-520c-3p | 1.4 | 0.00051 | 0.04135 |
| hsa-miR-424-3p | -1.9 | 0.00053 | 0.04135 |
| hsa-miR-135a-5p | 2.1 | 0.00058 | 0.04135 |
| hsa-miR-518f-3p | 1.3 | 0.00075 | 0.04680 |
| hsa-miR-519a-3p | 1.4 | 0.00079 | 0.04680 |
| hsa-miR-519b-3p | 1.6 | 0.00094 | 0.04878 |
| hsa-miR-512-3p | 1.1 | 0.00097 | 0.04878 |

Low VO<sub>2</sub>peak (12.8±1.1 mL/kg/min). High VO<sub>2</sub>peak (33.4±4.0 mL/kg/min).

### Sex Differences in Expression of Skeletal Muscle MiRNAs and Genes

The expression of 25 miRNAs significantly differed between men and women participants. Among the 25 miRNAs, 14 remained significant after p-value adjustment (**Table 3**). Interestingly, the expression of 8 of the 14 miRNAs showed significant differential expression between low and high VO_2_peak (miR-520g-3p, miR-520c-3p, miR-519c-3p, miR-517a-3p, miR-517b-3p, miR-518f-3p, miR-515-5p, and miR-503-5p), thus exhibiting both sex and VO_2_peak differences. In addition, the expression of 3,032 genes significantly differed between men and women and 917 genes remained significant after p-value adjustment (**Table S3**).

**Table 3:**
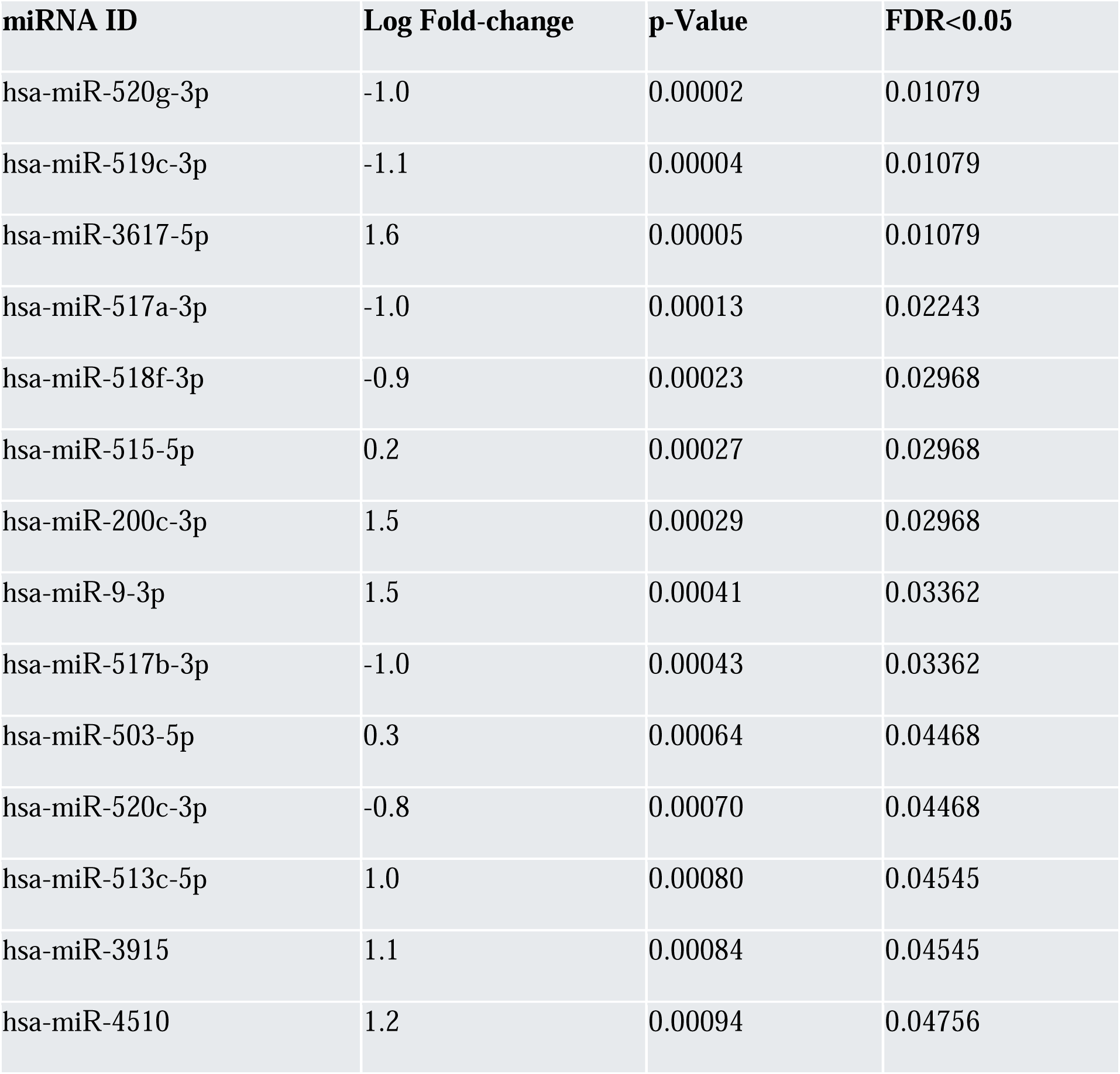
Differentially expressed skeletal muscle miRNAs between men and women VO_2_peak participants.

### Expressed MiRNAs and Genes that Correlate with VO**_2_**peak

We used LIMMA software for regression analysis of validation group samples and observed that 100 miRNAs showed significant correlation with VO_2_peak after p-value adjustment, including miR-503-5p and miR-206 (**Table S4**). After age/sex adjustment, 21 miRNAs remained significantly correlated with VO_2_peak (r ≥ 0.3; FDR adjusted p < 0.05). Among the 21 miRNAs identified, 14 were differentially expressed between low and high VO_2_peak participants in the discovery group. In addition, the expression of 3,962 genes significantly correlated with VO_2_peak, and 1,744 genes remained significant after age/sex adjustment (r ≥ 0.3; FDR adjusted p < 0.05) (**Table S5**). Among the 1,744 genes, 1,290 were differentially expressed between low and high VO_2_peak participants.

### MiRNA Target Genes and Molecular Networks Related to VO_2_peak

Of the 2,018 genes that significantly differed between low and high VO_2_peak individuals after p-value adjustment, 142 are regulated by the 14 differentially expressed miRNAs. Of these 14 miRNAs, the expression of 10 miRNAs (miR-519a-3p, miR-520g-3p, miR-515-5p, miR-517-3p, miR-518-3p, miR-512-3p, miR-424-3p, miR-135a-5p, miR-503-5p, and miR-291a-3p) showed inverse correlation with 50 target genes, which is consistent with miRNA functionality and role in regulating gene expression. **Figure 1** depicts the miRNA-gene molecular network consisting of the differentially expressed miRNAs and their target genes. **Figure 2A** shows the pathways enriched with miRNA target genes, including mitochondrial ATP synthesis and dysfunction, oxidative phosphorylation and TCA cycle, sirtuin1, and endothelial nitric oxide (NO) signaling pathways. **Figure 2B** depicts the proportions of genes upregulated or downregulated in these pathways.

**Figure 1:**
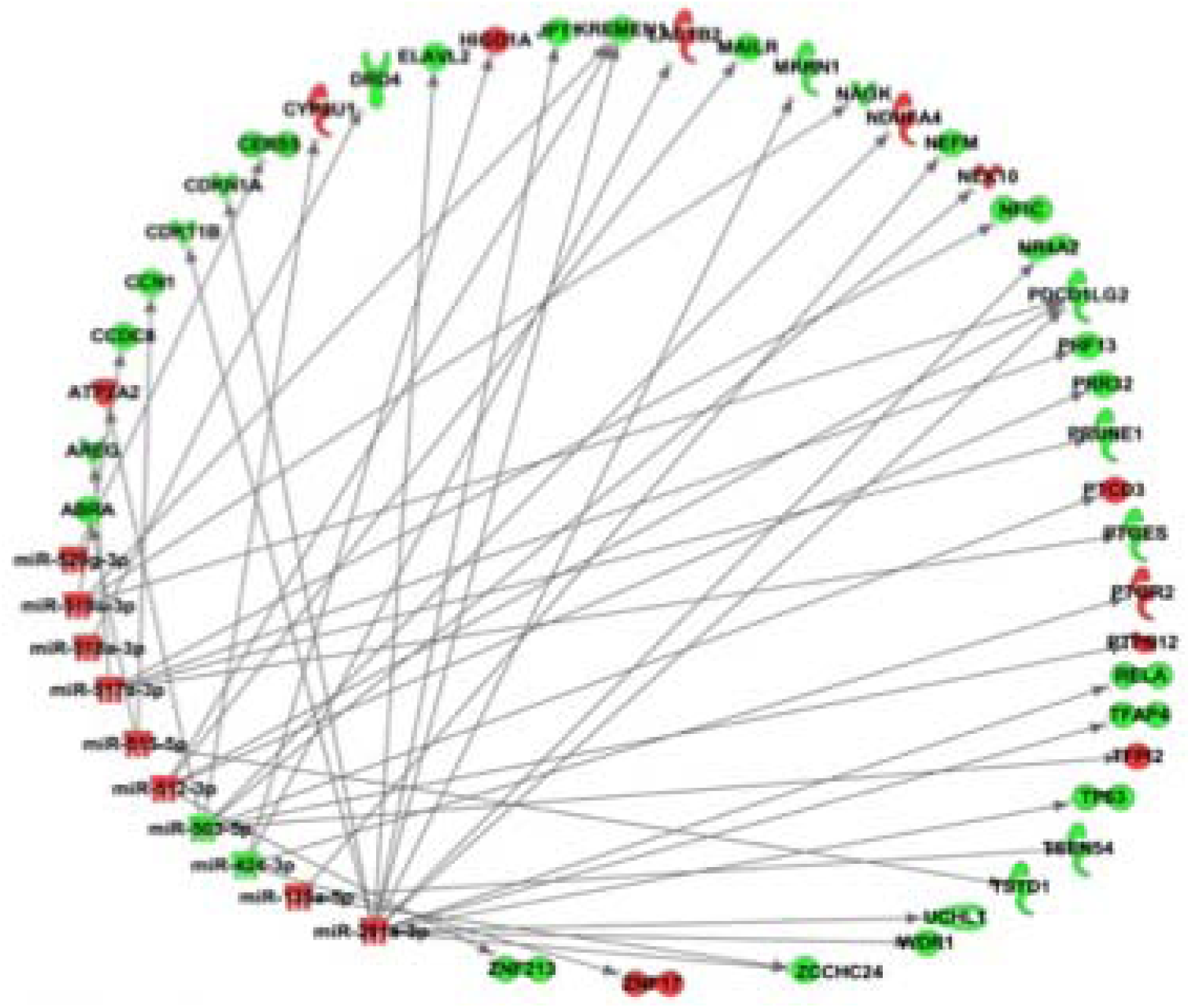
Skeletal muscle miRNA-gene network associated with VO_2_peak. The expression of miRNAs and miRNA target genes was analyzed from same skeletal muscle samples using untargeted sequencing. The expression datasets of miRNAs and genes were integrated using Ingenuity Pathway Analysis tool to identify the miRNA-gene network. miRNAs and miRNA target genes are represented as nodes. Red and green nodes indicate upregulated and downregulated miRNAs and genes. The molecular relationships between nodes are presented as a line (edge); arrows indicate the direction

**Figure 2.**
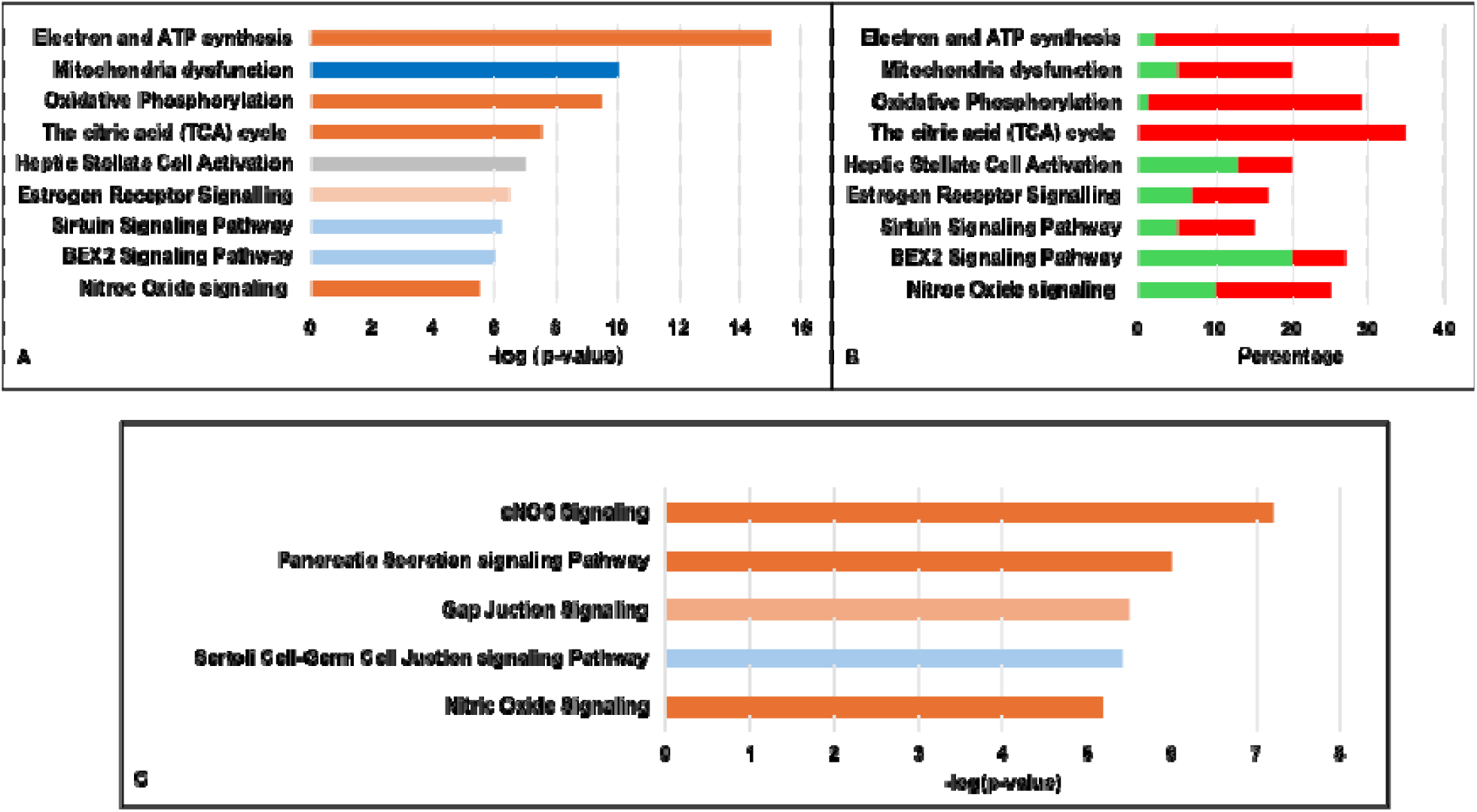
Enrichment Analyses. We used IPA (QIAGEN) and embedded miRNA target predication algorithms to identify inversely correlated expressed miRNAs and miRNA target genes. Core analysis tool embedded in IPA was used to identify enriched pathways. **(A)** Pathways enriched with miRNA target genes. Solid brown color indicates activated pathways and blue color indicate inhibited pathways. Horizontal axis depicts -log (p-value), with vertical line showing significant statistical threshold. **(B)** Proportion of genes upregulated or downregulated in the pathways enriched with miRNA target genes. Red color indicates upregulated genes, and blue color indicates downregulated genes in a pathway. The number of genes involved in a pathway is shown on the extreme right. **(C)** Pathways enriched with miRNA target genes in the MM Cyan module. WGCNA findings identified MM Cyan module as the top gene module that significantly correlated with VO_2_peak. Solid brown color indicates activated pathways and blue color indicate inhibited pathways. Horizontal axis depicts -log (p-value), with vertical line showing significant statistical threshold.

### Gene Modules Correlated with VO_2_peak and Enriched Pathways

For the differentially expressed genes predicted to be regulated by the 14 miRNAs, we performed WGCNA, an unbiased method to construct molecular networks based on pairwise correlations between variables to identify gene modules that significantly correlate with VO_2_peak variation. We identified 10 modules with r ≥ 0.30 and p-value < 0.05 (**Figure 3**). Among the 10 modules, ME Cyan module showed the highest correlation and significant level (r=0.76 and p=4×10^-12^). We identified significant pathways enriched with the ME Cyan module genes, including endothelial NO signaling (**Figure 2C**), which is relevant to cardiovascular system and cardiorespiratory fitness.

**Figure 3:**
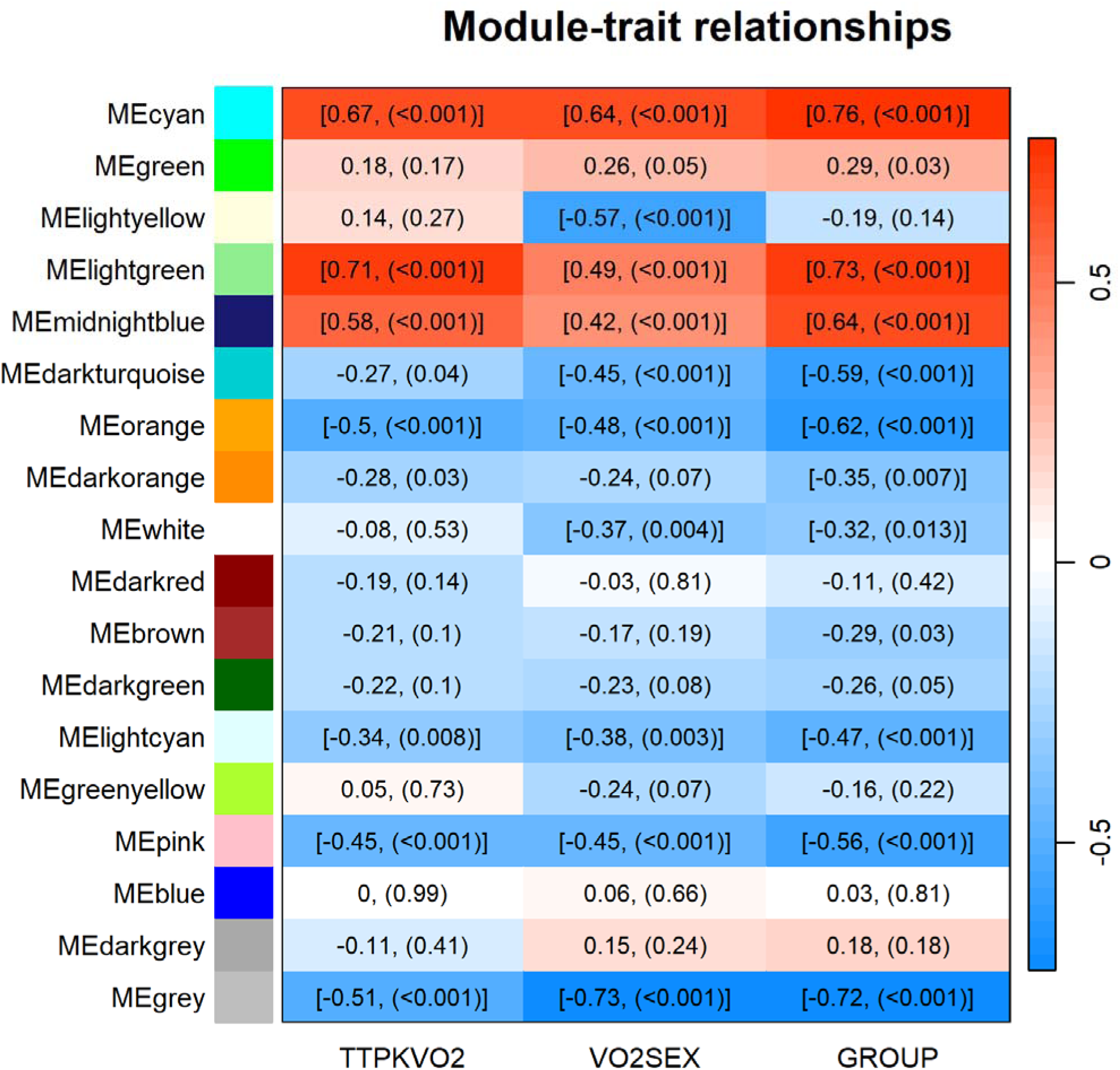
Correlation matrix showing gene modules that correlate with VO_2_peak in SOMMA participants. Each row corresponds to a module, bottom columns represent different traits: VO_2_peak for all participants (TTPKVO2), VO_2_peak dichotomized in men and women (VO2SEX) and VO_2_peak dichotomized in low and high participants (GROUP). Numbers in each row indicates correlation r-values and p-values in parenthesis. Brackets indicate modules we considered significantly correlated with a trait.

## DISCUSSION

This is the first study to identify a network of genes that are differentially regulated by a pool of miRNAs distinctly associated with VO_2_peak in older adults. We leveraged baseline skeletal muscle samples from a cohort of well-characterized SOMMA participants (30). We have identified expressed miRNAs and genes that correlate with and differ between low and high VO_2_peak older adults and show sex differences. Further, we have identified miRNA target genes, molecular networks, and enriched pathways related to VO_2_peak variation. These findings may provide novel insights into potential mechanisms that may drive skeletal muscle aging and decline in VO_2_peak among older adults, and potential therapeutic targets to slow skeletal muscle aging to mitigate impaired mobility and reduce mortality of older adults.

### Expressed MiRNAs and Genes

Age-associated dysfunction of skeletal muscle is associated with morbidity, mortality, and overall decline of quality of life in the older adults (1, 2). With aging, there is a decline in muscle mass, strength, and aerobic capacity, consistent with lower VO_2_peak (5–9). The etiology of lower VO_2_peak with aging is also partially due to lower maximal heart rate and thus cardiac output (1, 2). To our knowledge, no prior study compared expressed skeletal muscle miRNAs between individuals with low and high VO_2_peak. We report 14 novel miRNAs that significantly differ between individuals with low and high VO_2_peak and correlate with VO_2_peak after sex/age and p-value adjustments. Findings from our previous study have shown that 7 of the 14 miRNAs are also differentially expressed in serum samples from the same SOMMA cohort (38).

Despite the limited knowledge on the skeletal muscle miRNAs that may regulate VO_2_peak homeostasis, accumulating evidence demonstrates that miRNAs expression is dysregulated in response to skeletal muscle aging in mice, rhesus macaque and humans (13, 26–28), including miR-542, miR-133a and let-7f. It is important to note that these studies did not apply multiple testing in the data analysis. Here we show in statistical analysis prior to p-value adjustment that these miRNAs significantly differ between low and high VO_2_peak individuals. Further, we observed that four (miR-1, miR-133a, miR-206, miR-208a/b) of the eight known striated muscle-specific expressed miRNAs (39) were differentially expressed between low and high VO_2_peak participants; miR-206 is exclusively expressed in skeletal muscles (39). Our findings and previous observations suggest that some miRNAs dysregulated in response human skeletal muscle aging could be involved in VO_2_peak variation associated with physical functionality.

Beyond miRNA expression, other studies have delineated genes expressed in human skeletal muscle related to aging, type 2 diabetes and dystrophy (40, 41). In a recent study, 11 genes (NEFM, CDH15, RUNX1, CHRNA1, CHRND, NCAM1, MUSK, CDK5R1, CAV3, SCN4A, and SCN5A) related to skeletal muscle denervation were negatively associated with VO_2_peak and mitochondrial respiration (32). In our study, the expression of all 11 genes was significantly decreased in individuals with high VO_2_peak compared to participants with low VO_2_peak after p-value adjustment (FDR adjusted p <0.05). Further, our study confirmed significant negative correlation with VO_2_peak for 7 of the 11 genes (NEFM, CDH15, RUNX1, CHRNA1, CAV3, SCN4A, and SCN5A) after p-value and age/sex adjustments. These findings highlight the significance of denervation genes in skeletal muscle aging and VO_2_peak variation in older adults.

### Molecular Significance of the Identified MiRNA-Gene Molecular Network

Despite the advancement in knowledge on miRNAs and genes expressed in aging skeletal muscle, the molecular mechanisms that link skeletal muscle aging and decline in VO_2_peak remain poorly understood. To fully understand such mechanisms, it is necessary to interrogate miRNAs and genes expressed in same skeletal muscle samples derived from older adults discordant for VO_2_peaks. Further, it is important to assess expressed miRNAs and genes using untargeted and sensitive approaches such as high throughput sequencing to enable identification of novel and low abundant molecules. SOMMA is the ideal study to answer questions related to the molecular mechanisms responsible for decline in VO_2_peak in older adults because of the availability of extensive clinical data and skeletal muscle biopsies. Therefore, we integrated the datasets of the differentially expressed miRNAs and the differentially expressed genes to identify miRNA target genes. We identified a molecular network of 10 miRNAs that show inverse correlation with and regulate 50 target genes. These key miRNAs included a cluster of miR-519a/b/c, which was significantly increased in individuals with high VO_2_peak compared to subjects with low VO_2_peak. Interestingly, we previously reported that miR-519a-3p is significantly elevated in serum samples from the same SOMMA cohort (38). To our knowledge, no prior study has associated miR-519a-3p with skeletal muscle aging and/or VO_2_peak; however, there is cumulative evidence that the expression of miR-519a-3p is related with cellular processes including proliferation and migration and in various cancer types (30, 31, 32). Further interrogation revealed potential target genes regulated by miR-519a-3p, including MCM7 and PDCD1LG2, which are involved in the regulation of cellular processes related to cancer development (42, 43). The aberrant expression of miR-519a-3p in skeletal muscles and in serum from same individuals and the association with VO_2_peak variation suggest a novel biomarker and potential therapeutic target to slow skeletal muscle aging and elevate VO_2_peak.

Other remarkable novel miRNA-gene pair expressed in skeletal muscles include miR-503-5p-NDUFA4 axis. miR-503-5p was significantly decreased in high VO_2_peak individuals relative to low VO_2_peak study participants. We also identified miR-503-5p in serum skeletal muscle-specific exosomes (unpublished data), suggesting a potential molecular biomarker, cell to cell communication mediator as well as a potential therapeutic target to slow skeletal muscle aging. Further, we observed that miR-503-5p regulates NDUFA4, a key modulator of mitochondrial respiration (44). NDUFA4 is a subunit of mitochondrial complex VI, cytochrome c oxidase, responsible for mitochondrial respiration. Mitochondrial dysfunction is related to VO_2_peak (8, 45) and our study suggests that miR-503-3p-NDUFA4 axis could be a potential regulator of skeletal muscle function and VO_2_peak by modulating mitochondrial respiration.

Overall, the identified molecular network of miRNAs and genes may highlight the significance of skeletal muscle molecular changes in VO_2_peak variation in older adults. Further, the network provides opportunities for generation of hypotheses to test the significance of miRNA-mediated molecular mechanisms that underlie VO_2_peak variation.

### Enriched Pathways related to VO_2_peak

Our findings show that miRNAs may regulate key pathways enriched with miRNA target genes expressed in skeletal muscles to influence skeletal muscle aging and associated VO_2_peak trait. We used IPA tool to analyze a dataset of miRNA target genes and identify enriched pathways. We observed that mitochondrial activity (ATP synthesis, oxidative phosphorylation and TCA cycle), sirtuin, BEX2 signaling pathways as well as endothelial NO signaling are activated in high VO_2_peak compared to low VO_2_peak individuals. In contrast, mitochondria dysfunction was inactivated in individuals with high VO_2_peak. Likewise, we observed that a higher percentage of upregulated genes are involved in mitochondrial activity, sirtuin signaling, and nitric oxide signaling in participants with high VO_2_peak. Further analyses of miRNA target genes using WGCNA implemented in R package revealed 10 gene modules significantly correlated with VO_2_peak trait. A pathway enrichment analysis of the genes contained in the ME Cyan module, with the strongest correlation with VO_2_peak, confirmed that nitric oxide signaling is one of the top pathways related toVO_2_peak. These observations are consistent with the previous findings on the role of these pathways in skeletal muscle aging. For example, previous studies demonstrated that miRNAs regulate key genes in sirtuin1 signaling, which is involved in protection against cell senescence to slow skeletal muscle aging (46–48); decline in mitochondrial activity is associated with skeletal muscle aging, reduced VO_2_peak, walking speed, leg power, and recurrent falls (6, 8, 45, 49). Our observations indicate that key pathways detrimental to the health of skeletal muscles such as the mitochondrial dysfunction are inactivated in individuals with high VO_2_peak and activated in subjects with low VO_2_peak. Thus, potential miRNA therapeutics could reduce the effects detrimental pathways for individuals with low VO_2_peak to improve physical function.

## LIMITATIONS

The key limitations are the cross-sectional nature of the study, the unbalanced ethnic/racial composition of the study participants, and the lack of sex/ethnic stratified analysis that limits generalization. Although our analysis included sex and age as confounding factors, the findings are limited due to the small sample size, which constrained our ability to adjust for additional covariates. As a result, we could not account for other potentially confounding factors, including comorbidities, medications, physical activity levels, or diet that might influence miRNA expression and VO_2_peak. Thus, our results should be cautiously interpreted. Therefore, there is a need to validate the findings, adjusting for appropriate cofounding factors, in larger longitudinal cohorts of SOMMA participants.

## CONCLUSION

We identified miRNAs and miRNA target genes, coordinately regulated molecular networks, and pathways related to VO_2_peak variation, including mitochondrial respiration and NO signaling. Our findings highlight critical insights into the molecular changes that drive low VO_2_peak and potential therapeutic targets for intervention to slow skeletal muscle aging to reduce immobility and mortality in older individuals. Future studies will validate the findings in a larger, longitudinal study cohort, and interrogate, both in-vitro and in-vivo, the miRNA-mediated molecular mechanisms that underlie VO_2_peak variation.

## ACKNOWLEDGEMENT

The authors gratefully acknowledge the editorial assistance of Indra M. Newman, PhD (Wake Forest Clinical and Translational Science Institute, which is supported by the National Center for Advancing Translational Sciences (NCATS) and National Institutes of Health under award number UM1TR004929).

## AUTHORS’ CONTRIBUTIONS

GMK: Conceptualization; Funding acquisition; Investigation; Methodology; Supervision; Visualization; Writing original draft; Writing-review and editing.

FH: Formal analysis; Investigation; Methodology; Visualization; Writing-review and editing.

RH: Investigation; Writing-review and editing.

PMC: Investigation; Writing-review and editing.

SRC: Investigation; Funding Acquisition; Writing-review and editing.

ABN: Investigation; Writing-review and editing.

NWG: Investigation; Writing-review and editing.

LMS: Investigation; Writing-review and editing.

NEL: Investigation; Writing-review and editing.

JX: Formal analysis; Software; Writing-review and editing

NTW: Methodology; Writing-review and editing

GL: Methodology; Writing-review and editing

JEC: Formal analysis; Software; Writing-review and editing.

LAC: Investigation; Supervision; Resources; Writing-review and editing.

SBK: Conceptualization; Investigation; Resources; Supervision; Writing-review and editing.

